# *TaBln1* negatively regulates wheat resistance to stripe rust by reducing Ca^2+^ influx

**DOI:** 10.1101/2021.07.16.452683

**Authors:** Shuangyuan Guo, Yanqin Zhang, Peng Zeng, Min Li, Qiong Zhang, Xing Li, Quanle Xu, Tao Li, Xiaojie Wang, Zhensheng Kang, Xinmei Zhang

## Abstract

Blufensin1 (Bln1) has been identified as a negative regulator of basal defense mechanisms that is unique to the cereal grain crops barley, wheat, and rice. However, the molecular mechanisms through which *Blufensin1* regulates the wheat immune response are poorly understood. In this study, we found that *TaBln1* is significantly induced by *Puccinia striiformis* f. sp. *tritici* (*Pst*) virulent race CYR31 infection. Knockdown the expression of *TaBln1* by virus-induced gene silencing reduced *Pst* growth and development, and enhanced the host defense response. In addition, TaBln1 was found to physically interact with TaCaM3 on the plasma membrane. Silencing *TaCaM3* with virus-induced gene silencing increased fungal infection areas and sporulation and reduced wheat resistance to the *Pst* CYR23 and CYR31. Moreover, we found that the *TaCaM3* transcription level could be induced by treatment with chitin but not flg22. Silencing *TaCaM3* decreased the Ca^2+^ influx induced by chitin, but silencing *TaBln1* increased the Ca^2+^ influx *in vivo* using a non-invasive micro-test technique. Taken together, we identified the wheat negative regulator TaBln1, which interacts with TaCaM3 to impair Ca^2+^ influx and inhibits plant defenses.

**One-sentence summary:** *TaBln1* negatively regulate wheat resistance to stripe rust possibly due to the interaction with TaCaM3 on the plasma membrane, which impairs the calcium influx modulated by TaCaM3.

## Introduction

Negative regulators of plants, a type of susceptibility factor encoded by susceptibility genes (S genes), are recruited and utilized by pathogens to interfere with plants’ defensive responses; this directly or indirectly helps the growth and parasitism of pathogens (Pavan et al., 2010; Lapin et al., 2013; Schie et al., 2014). Altering S genes could result in a more broad-spectrum and durable natural resistance in plants (Pavan et al., 2010; Schie et al., 2014). Some successful examples are the use of *mildew-resistance locus O* (*MLO*) mutations in barley, wheat, and tomato, which confer heritable resistance to powdery mildew. Loss of function in *MLO* mutants results in durable and broad-spectrum resistance (Jørgensen, 1992; Acevedo-Garcia et al., 2015; Wang et al., 2014; Nekrasov et al., 2017). Likewise, simultaneous mutation of the three homologues of *enhanced disease resistance 1* in wheat using CRISPR/Cas has enhanced resistance to powdery mildew (Zhang et al., 2017). Some researchers have generated rice lines with broad-spectrum resistance to *X. oryzae* pv. *oryzae* by editing the promoter of three *SWEET* genes using CRISPR/Cas9 in rice (Oliva et al., 2019; Xu et al., 2019). Similarly, *Pi21* encodes a proline-rich protein in which mutation through RNA interference can confer resistance to *M. oryzae* in rice (Fukuoka et al., 2009). Another example involves RACB, a member of the plant RAC/ROP family of RHO-like small monomeric G-proteins. Transient RNA interference through the double-stranded RNA (dsRNAi) of barley *HvRacB* induces partial *Bgh* resistance (Schultheiss et al., 2002; 2003). These results clearly show that identifying the precise role of negative regulators in plants is part of a powerful approach for generating resistance in plants.

*Blufensin1* (*Bln1*), which is a member of a new small peptides family, was initially identified as a negative regulator of basal defense mechanisms (Caldo et al., 2004). Computational interrogation of the *Bln1* gene family determined that these peptides are uniquely encoded in the cereal grain crops barley, wheat, and rice. Suppressed expression of *Bln1* via barley stripe mosaic virus (BSMV)-induced gene silencing decreased susceptibility in compatible interactions between barley and powdery mildew. However, overexpression of *Bln1* significantly increased accessibility to virulent *Blumeria graminis* f. sp. *hordei* (*Bgh*) (Meng et al., 2009). Recently, transient overexpression of *Bln2* (a blufensin with high sequence similarity to *Bln1*) in barley has been found to increase the susceptibility of barley to *Bgh* as well. Interestingly, bimolecular fluorescence complementation (BiFC) assays have shown that BLN1 and BLN2 can interact with the Ca^2+^ sensor calmodulin (CaM) (Xu et al., 2015). However, despite these advances, it remains to be seen whether family members of BLN and CaM antagonize or cooperate with each other in modulating immune responses in wheat.

In plant cells, calcium ions are a ubiquitous intracellular second messenger involved in numerous signaling pathways, particularly in relation to the triggering of immune responses (Tuteja et al., 2007; Hilleary et al., 2020). Ca^2+^ elevation in the cytosol is an early essential event in pathogen response signaling cascades and leads to the activation of downstream innate immune responses (Dangl et al., 1996). For example, the recognition of the pathogen results in cyclic nucleotide production and the activation of cyclic nucleotide-gated channels, which leads to downstream generation of pivotal signaling molecules. CaMs and CaM-like proteins (CMLs) are involved in this signaling, functioning as Ca^2+^ sensors and mediating the synthesis of NO(Ma et al., 2011). CaMs and CMLs constitute a large family of Ca^2+^ binding and signaling proteins in plants. Among these proteins, CaM, containing a pair of Ca^2+^-binding EF-hand motifs, is the most thoroughly characterized Ca^2+^ sensor in plants (Bouché et al., 2005; Defalco et al., 2009). Plant genomes contain multiple loci that encode conserved CaM isoforms. For example, seven distinct genes encode four protein isoforms (CaM2/CaM3/ CaM5, CaM1/CaM4, CaM6, and CaM7) in *Arabidopsis* (McCormack et al., 2003). It has been reported that CaMs are involved in innate immunity in plants, which can transmit the initial signal of cytosolic Ca^2+^ elevation to downstream targets in signal transduction cascade (Ma et al., 2007). Pathogen infection results in induction and/or suppression of plant CaM isoforms (Garcia-Brugger et al., 2006). Manipulating plant CaM expression affects basal resistance to a range of pathogens (Heo et al., 1999; Takabatake et al., 2007). For example, the silencing of specific pathogen-induced CaM isoforms in tomato results in enhanced susceptibility to virulent necrotrophic bacteria and fungi (Takabatake et al., 2007). The expressions of *SCaM4* and *SCaM5* in transgenic tobacco and *Arabidopsis* lead to increased pathogenesis-related gene expression and enhanced resistance to bacterial, fungal, and viral pathogens (Zhu et al., 2010). CaM may be a key player in transducing pathogen-induced Ca^2+^ increase to the downstream components of defense signaling and contributing to plant defense responses.

Wheat (*Triticum aestivum* L.) is one of the most important dietary crops in the world. While growing, wheat undergoes continuous exposure to abiotic and biotic stresses in the natural environment. *Puccinia striiformis* f. sp. *tritici* (*Pst*), an obligate biotrophic pathogen, is the causal agent of wheat stripe rust and has become the largest biotic limitation to wheat production and the global food supply (Liu et al., 2016; Schwessinger, 2017). Genetic control is the most cost-effective strategy for reducing the threat of this disease (Ellis et al., 2014). Recent studies have indicated that broad-spectrum resistance can result from loss of function or a mutation of negative regulators (Qiang et al., 2021). Therefore, a better understanding of the mechanisms of this pathogen at the molecular level is of great importance to improve disease resistance in wheat.

We identified and functionally characterized one *Blufensin* gene, *TaBln1*, and assessed its important role in wheat resistance to *Pst*. In addition, we demonstrated that TaBln1 interacts with one CaM, TaCam3, which was shown to play a positive role in resistance to stripe rust in wheat. Concurrently, a noninvasive micro-test technique (NMT) and chitin-treatment experiments showed that transient silencing of *TaCaM3* led to a decrease in calcium influx in wheat mesophyll cells; however, silencing *TaBln1* led to a significant increase in calcium influx. Our results demonstrate that *TaBln1* negatively regulates wheat resistance to *Pst* by affecting the balance between TaCam3 and calcium, and TaCam3 might be the target for TaBln1 in the wheat response to *Pst* infection. Our results provide new insights that may lead to improved understanding of the roles of negative regulators in plant resistance.

## Results

### *TaBln1* is significantly induced by the *Pst* virulent race CYR31

Using a complete open reading frame, the novel gene *TaBln1* was isolated from the transcriptome of wheat cultivar Suwon 11 infected by virulent *Pst* CYR31. BLAST analyses found in the Ensemble Plants database revealed that three copies of *TaBln1* were located on chromosomes 4A, 5B, and 5D, but no complete sequence information was found. Subsequently, the copies of *TaBln1* were obtained by genomic PCR with specific primers (Supplemental Table 1), according to the wheat genome database. The three copies shared 97.33% DNA identity and 97.56% protein identity, indicating that they may possess identical biological functions.

To investigate whether *TaBln*1 participates in the wheat response to *Pst*, we used RT-qPCR to analyze the relative transcript profiles of *TaBln* with a pair of conservative primers in the *Pst* virulent race CYR31 and the avirulent race CYR23 in inoculated wheat at different time points. The expression of *TaBln1* was upregulated at 12, 24, and 72 h post inoculation (hpi) in wheat leaves challenged with the *Pst* virulent race CYR31 (compatible interaction). The highest *TaBln1* transcript level was approximately 9-fold at 24 hpi. However, the expression of *TaBln1* showed no significant change in wheat leaves challenged with the avirulent race CYR23 (Figure 1), suggesting that *TaBln1* may participate in a compatible interaction between wheat and *Pst*.

**Figure 1.**
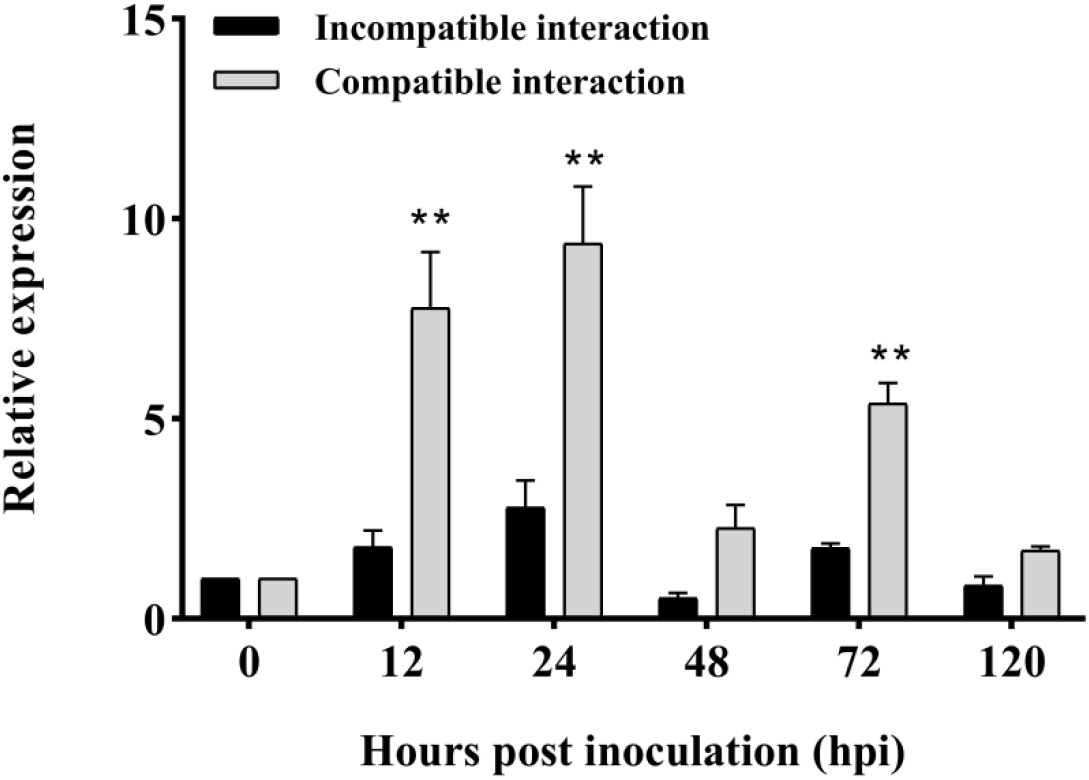
*TaBln1* is significantly induced by *Pst* race CYR31 infection. Wheat leaves were sampled for CYR23 and CYR31-inoculated plants at 0, 12, 24, 48, 72, and 120 hpi. Untreated leaves act as the control. The relative expression of *TaBln1* was calculated using the comparative threshold method (2^-ΔΔCT^). Expression levels were normalized to *TaEF-1a*. Values were derived from three biological repetitions. Asterisks indicate a significant difference (**P < 0.01) compared to the untreated control according to Student’s *t* test.

### Silencing of *TaBln1* reduces wheat susceptibility to *Pst*

To determine the role of *TaBln1* in the wheat–*Pst* interaction, we used a BSMV-induced gene silencing (VIGS) system to knock down the expression of *TaBln1*. Photobleaching was observed in plants inoculated with BSMV:TaPDS, and mild chlorotic mosaic symptoms were displayed in other BSMV-infected plants with no evident defects (Figure 2A). BSMV:TaPDS was used as a positive control to indicate the efficacy of VIGS system. Thereafter, the *Pst* CYR31 was used to inoculate the BSMV-treated plants. The disease phenotype showed that there were fewer uredinia on *TaBln1*-knockdown plants at 14 days post infection (dpi) than on the control plants (Figure 2B). Moreover, the fungal biomass was significantly lower on *TaBln1*-knockdown plants at 5, 7, and 10 dpi than on the control plants, which is consistent with the disease phenotype (Figure 2C). Standard curves (Supplemental Figure S2) for biomass were generated using qPCR with total genomic DNA (Yin et al., 2009). The transcriptional levels of *TaBln1* in BSMV:TaBln1-inoculated leaves were about 40–95% lower at 0, 24, 48, and 120 hpi than in control plants (Figure 2D), indicating that the expression of *TaBln1* was substantially knocked down by VIGS.

**Figure 2.**
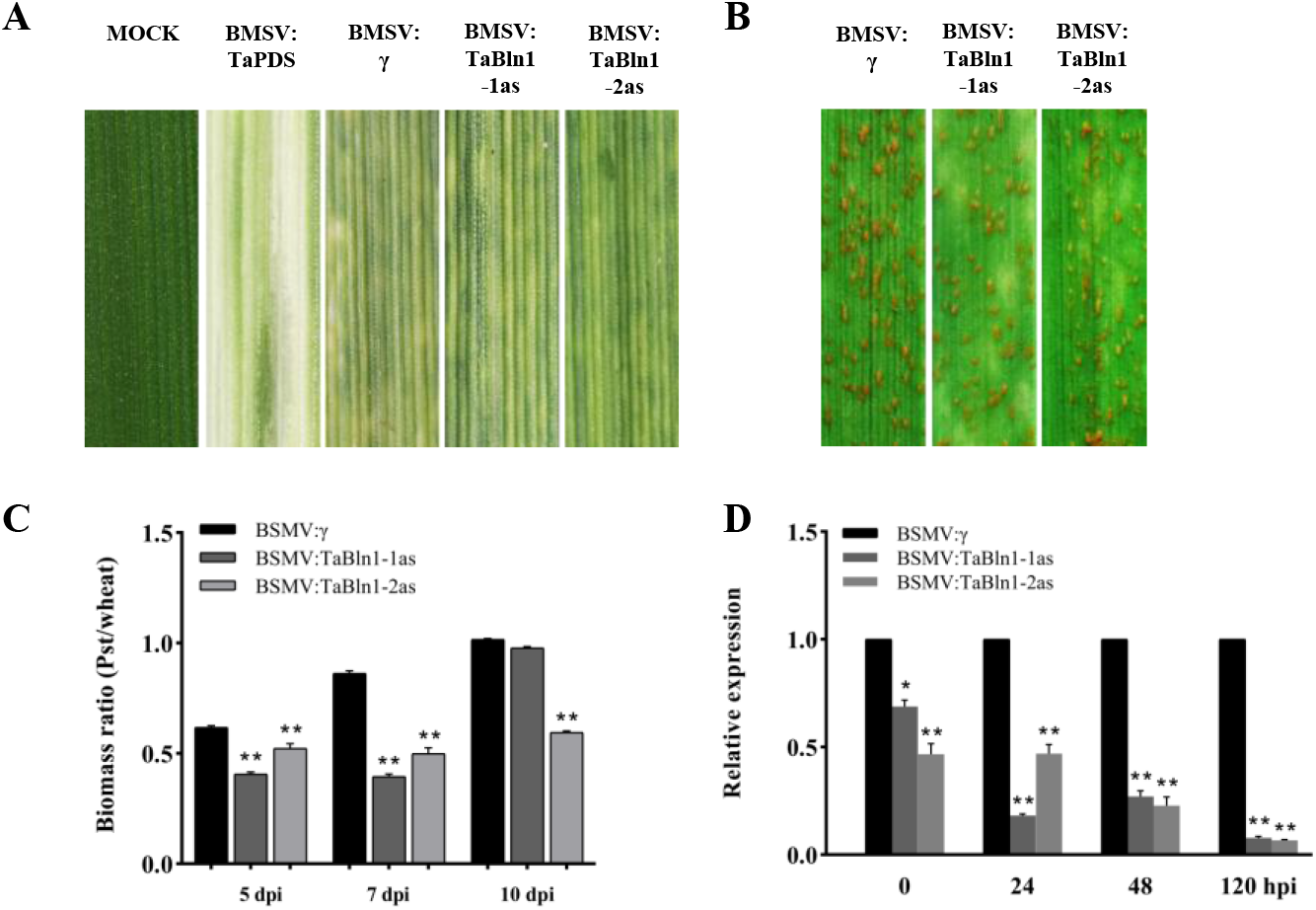
Silencing *TaBln1* reduces wheat susceptibility to *Pst* race CYR31. A, The phenotype did not change in wheat leaves mock inoculated. Photobleaching was visible on wheat leaves inoculated with BSMV:TaPDS. Slight chlorotic mosaic symptoms were observed in wheat leaves infected with BSMV:γ, BSMV:TaBln1-1as and BSMV:TaBln1-2as. B, Disease phenotypes of the fourth leaves pre-inoculated with BSMV constructs and then challenged with *Pst* CYR31. Disease phenotypes were photographed at 14 dpi. C, Biomass ratio of the *Pst*/wheat was assayed by qRT-PCR with total DNA extracted from BSMV-treated wheat leaves infected by CYR31 at 5, 7 and 10 dpi. Ratio of total fungal DNA to total wheat DNA was assessed by normalizing the data to the wheat gene *TaEF-1α* and the *Pst* gene *PstEF1*. D, The relative expression levels of *TaBln1* in wheat leaves inoculated with CYR31 were assayed by qRT-PCR at 0, 24, 48 and 120 hpi.

### Knockdown of *TaBln1* expression reduces *Pst* growth and development and enhances the host defense response

To quantify the reduced disease phenotype in *TaBln1*-knockdown plants, we also microscopically examined fungal growth and development in *TaBln1*-silenced leaves (Figure 3A). The number of hyphal branches and haustoria at 24 hpi and the length of hyphae at 48 hpi were significantly reduced in *TaBln1*-silenced plants relative to control plants inoculated with BSMV:γ (Figure 3B and C). Compared to the control plants, the infection areas of *Pst* at 120 hpi also decreased by silencing of *TaBln1* (Figure 3D). Moreover, to further analyze the host response, we measured the accumulation of H_2_O_2_ in *TaBln1*-knockdown plants with DAB staining. Our results revealed that the area that contained H_2_O_2_ was significantly larger in *TaBln1*-silenced leaves than in the control leaves at 24, 48, and 120 hpi (Supplemental Figure S3). These histological results indicate that the silencing of *TaBln1* restricted the growth and development of *Pst* and enhanced the resistance of wheat to *Pst*.

**Figure 3.**
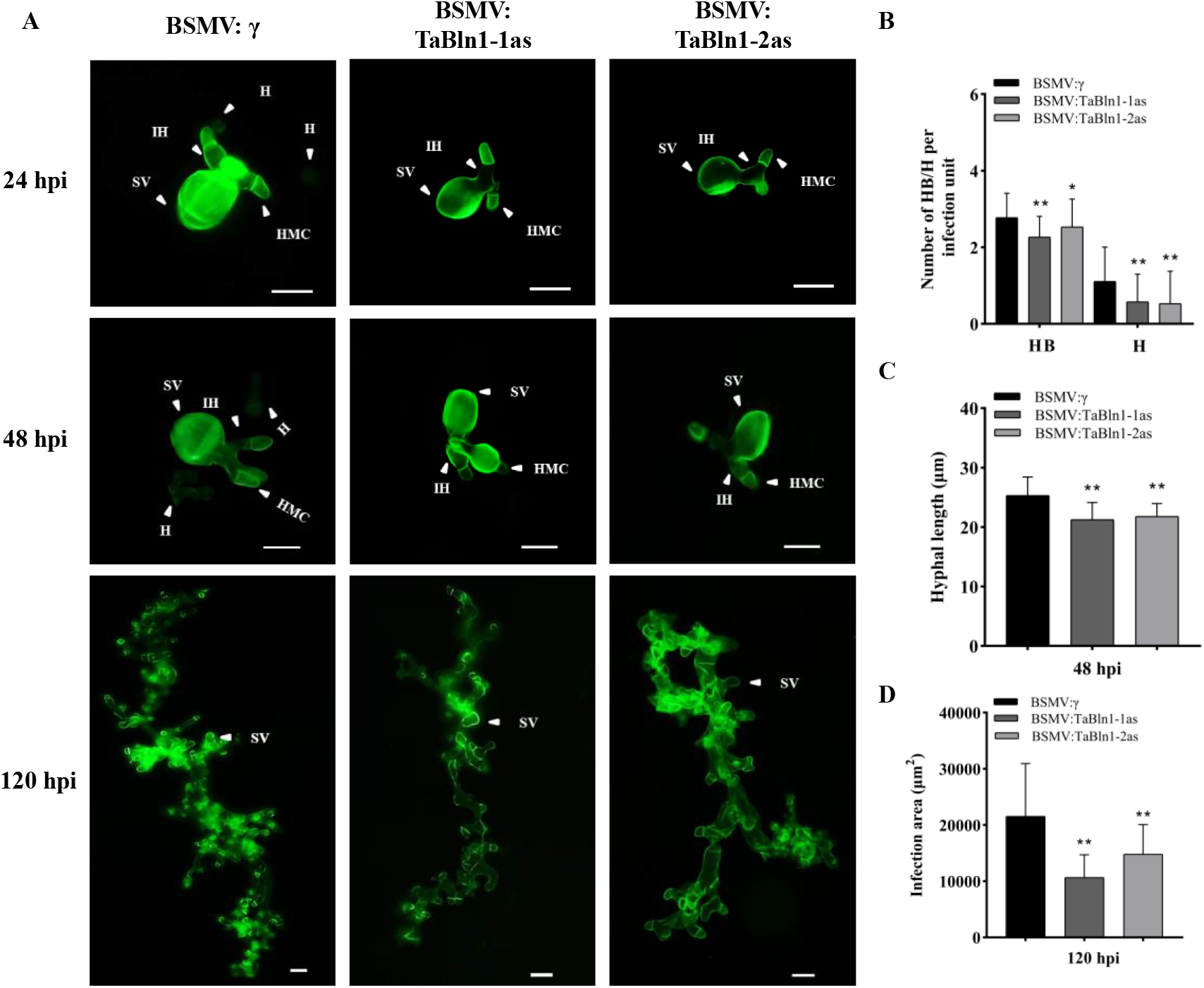
Knocking down *TaBln1* reduces growth and development of *Pst* race CYR31. A, Fungal structures in wheat leaves infected with BSMV and *Pst* were visualized with WGA and the fungal structures were observed under a fluorescence microscope. SV, substomatal vesicle; HMC, haustorial mother cell; IH, infection hypha. H, haustoria. B, The average number of hyphal branches (HB) and haustoria (H) of *Pst* in each infection site were counted at 24 hpi. C, The hyphal length of *Pst* in *TaBln1*-silenced plants at 48 hpi. Hyphal length is the average distance to the tip of the hypha from the intersection of the sub-stomatal vesicle and the hypha was measured using CellSens Entry software (unit in μm). D, The infection area at 120 hpi was calculated using DP-BSW software (units of μm^2^). Values were derived from three biological repetitions (50 infection sites each time). Asterisks indicate a significant difference (*P < 0.05, **P < 0.01) from BSMV:γ inoculated plants using the Student’s *t* test.

### TaBln1 interacts with the TaCaM3

BLN1 and BLN2 were found to interact with a CaM in barley (Xu et al., 2015), suggesting that TaBln1 may interact with a CaM in wheat. To confirm this prediction, *TaCaM3*, a homologue of barley CaM, was cloned in wheat cultivar Suwan11. BIFC was first used to detect whether TaBln1 interacts with TaCaM3 in transiently transformed *N. benthamiana* leaves. Strong fluorescence signals were obtained in the interaction between TaBln1 and TaCaM3 after 48 hpi (Figure 4A). However, there were no fluorescence signals for the co-expression of NE-TaBln1 and the empty vector CE in *N. benthamiana* leaves (Figure 4A). To further investigate their physical association with co-immunoprecipitation (Co-IP) assays, we transiently co-expressed TaBln1 fused with the GFP tag and TaCaM3 fused with the HA tag in *N. benthamiana*. As expected, TaCaM3 was co-immunoprecipitated with TaBln1 but not with GFP (Figure 4B). Next, their interaction was further confirmed in split-luciferase assays. The nLUC fusion of TaBln1 and cLUC fusion of TaCaM3 were co-expressed in *N. benthamiana*. Only the area co-expressing TaBln1-nLUC and TaCaM3-cLUC had a strong luminescence signal; no luminescence signals appeared in the area co-expressing TaBLN1-nLUC and cLUC, nLUC and TaCaM3-cLUC, or nLUC and cLUC (Figure 4C). In addition, the *in vitro* interaction between TaBln1 and TaCaM3 was tested using a pulldown assay with TaBln1 fused with GST tag and TaCaM3 fused with His tag. When detected with anti-His monoclonal antibodies, TaCaM3 was detected in Western blots of proteins eluted from TaBln1-GST beads, but no TaCaM3 was detected from free GST beads (Figure 4D). These data confirmed that TaBln1 interacts with TaCaM3.

**Figure 4.**
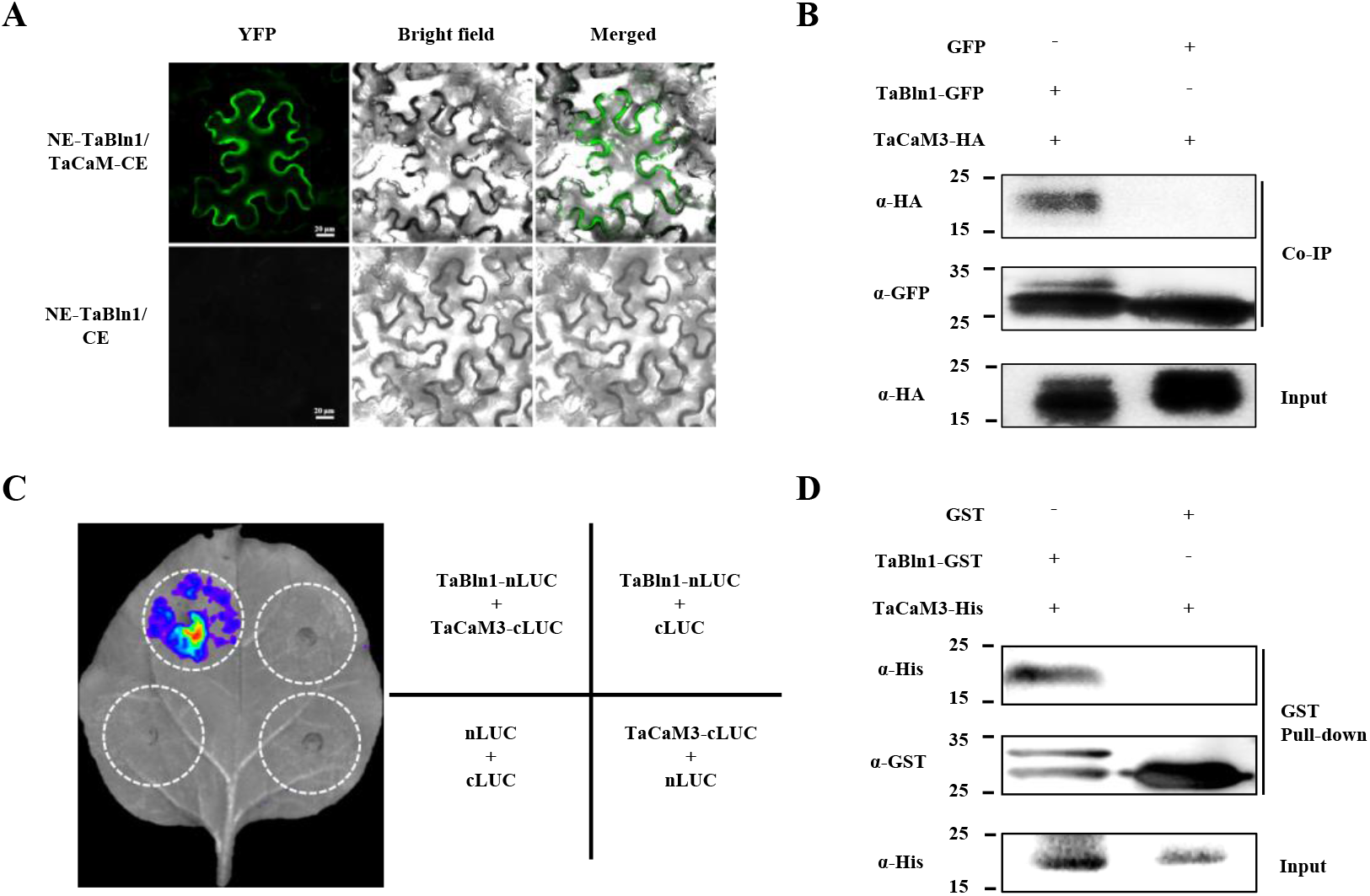
TaBln1 interacts with TaCaM3. A, BIFC confirmed the interaction of TaBlin1 and TaCaM3. NE-TaBlin1/TaCaM-CE is shown in the first panels, and NE-TaBlin1+CE as negative control is shown in the second panels. Bright-field and YFP fluorescence (in green) images were taken by confocal microscopy. Bar = 20 μm. B, Protein interaction of TaBln1-GFP and TaCaM3-HA were determined by Co-IP assay. Anti-HA and anti-GFP were used to detect protein expression. GFP was used as a negative control. kDa, kilodaltons. C, Split-luciferase assays demonstrated the interaction of TaBln1 and TaCaM3 in *N. benthamiana* leaves. The nLUC and cLUC alone were used as negative controls. D, GST pull down assay was used to detect the interaction between TaBln1 and TaCaM3. The recombinant TaBln1-GST and TaCaM3-His expressed in *E. coli* were subjected to GST pull down analysis. GST was used as a negative control. kDa, kilodaltons.

### TaCaM3 co-localizes with TaBln1 to the plasma membrane

To determine the subcellular localization of TaBln1, we first analyzed the protein sequence for TaBln1. TaBln1 contained a signal peptide (1–29 aa) and a putative transmembrane domain (Supplemental Figure S4), indicating that it might be localized to the biological membrane. To confirm this prediction, TaBln1-GFP and a marker protein of plasma membrane, TaWpi6-mCherry, were co-expressed in *N. benthamiana* leaves. As expected, GFP fluorescent signals of TaBln1-GFP were detected in the plasma membrane and nuclear membrane; meanwhile, the green fluorescence substantially overlapped with the red fluorescence of TaWpi6-mCherry (Figure 5A). To further validate the localization of TaBln1, plasmolysis was induced by the addition of 800 mM mannitol, and then, clear plasma membrane and nuclear membrane signals were observed for TaBln1-GFP (Figure 5B), indicating that TaBln1 was localized to the plasma membrane and nuclear membrane. In addition, in view of the interaction between TaBln1 and TaCaM3, we used TaCaM3-GFP and TaBln1-mCherry to determine whether TaBln1 co-localized with TaCaM3 to the plasma membrane and nuclear membrane. We determined the subcellular localization of TaCaM3, which, similar to the free GFP, was detected throughout the cytosol, plasma membrane, and nucleus (Supplemental Figure S5). However, the results of co-localization showed that the red fluorescence of TaBln1-mCherry substantially overlapped with the green fluorescence of TaCaM3-GFP in the plasma membrane but not with the nuclear membrane (Figure 5C). This suggests that TaBln1 mainly co-localized with TaCaM3 to the plasma membrane.

**Figure 5.**
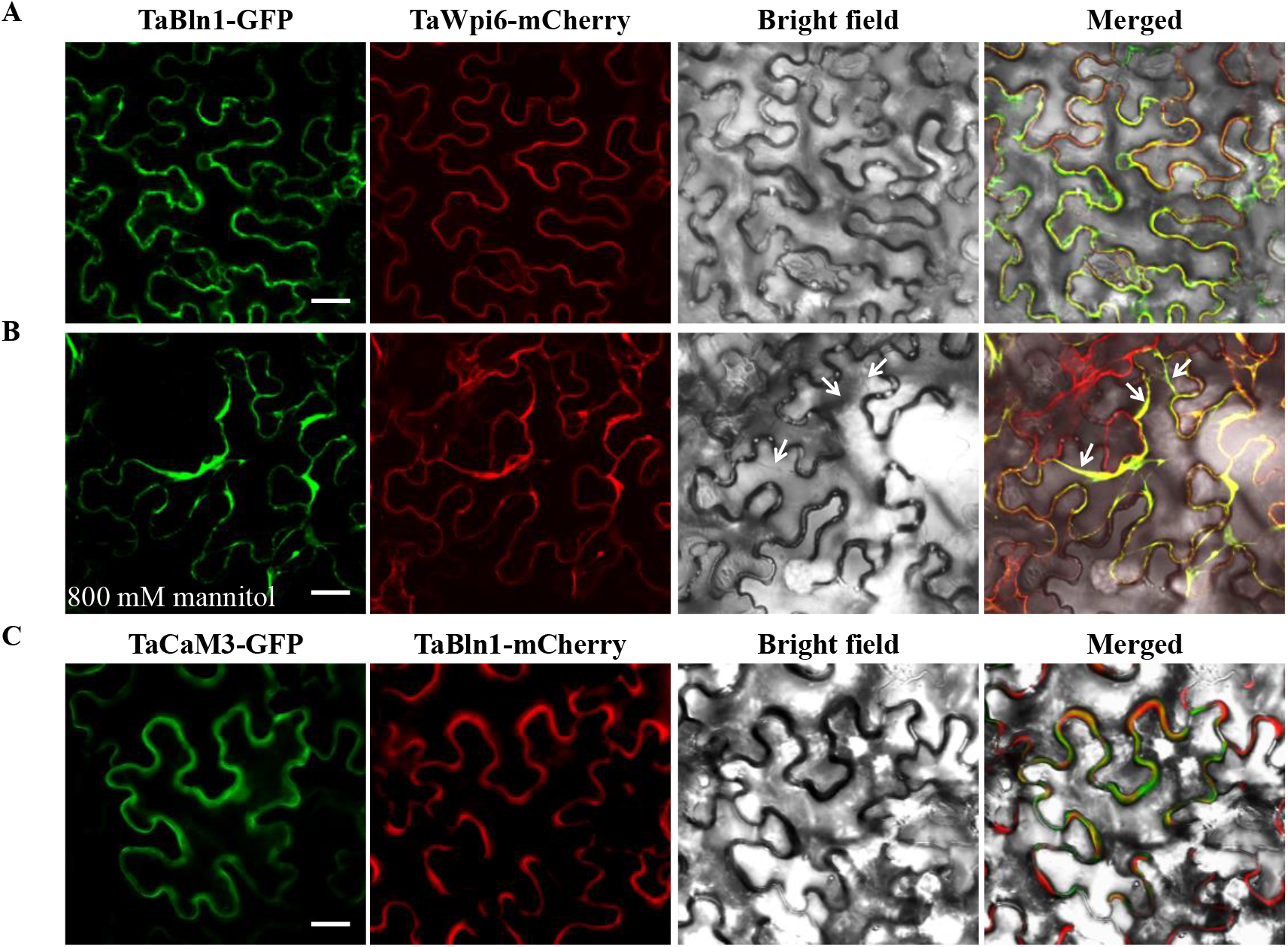
Subcellular localization of TaBln1 and co-localization of TaBln1 and TaCaM3 in *N. benthamiana* leaves. A and B, Co-expression of TaBln1-GFP and TaWpi6-mCherry in *N. benthamiana* leaves. GFP fluorescence is in green; TaWpi6-mCherry (red fluorescence) indicates TaWpi6 labeling the plasma membrane. Leaf epidermal peels in (B) were plasmolyzed (white arrows) by incubation in 800 mM mannitol for 6 min. C, TaCaM3-GFP and TaBln1-mCherry were co-expressed in *N. benthamiana* leaves. Bright field (BF) images show the equivalent field observed under white light. Merged TaBln1-GFP/TaWpi6-mCherry/Bright field (A and B) and TaCaM3-GFP/TaBln1-mCherry/Bright field (C) images are shown. All of the signals were monitored by confocal microscopy. Bar = 20 μm.

### Silencing of *TaCaM3* reduces wheat resistance to *Pst*

To obtain direct evidence for the function of *TaCaM3*, we first assayed the expression of *TaCaM3* at different levels of *Pst* infection. The six copies of *TaCaM3* on chromosomes 2A, 2B, 2D, 4A, 4B, and 4D share 95.5% similarity in nucleotide sequence (Supplemental Figure S6) and nearly 100% similarity in amino acid sequence (Supplemental Figure S7), which led to the design of conservative *TaCaM3* primers for RT-qPCR. The transcription of *TaCaM3* was significantly induced in both compatible and incompatible interactions at 12 and 24 hpi. The highest *TaCaM3* transcript level was approximately 3-fold in incompatible interaction at 48 hpi (Supplemental Figure S8), suggesting that *TaCaM3* plays a role in the interaction between wheat and *Pst*.

To determine the function of *TaCaM3* in the wheat–*Pst* interaction, we silenced *TaCaM3* using VIGS. When *TaCaM3*-knockdown plants were inoculated with CYR23, conspicuous HR was elicited on leaves that had previously been infected with BSMV:γ and BSMV:TaCaM3 (Figure 6A). However, a slight sporulation of *Pst* emerged around the necrotic regions on leaves infected with BSMV:TaCaM3 at 14 dpi (Figure 6A). Although there was no obvious difference between the BSMV:TaCaM3 and BSMV:γ plants in fungal biomass at 10 dpi, the fungal biomass was significantly increased in BSMV:TaCaM3 leaves at 5 and 7 dpi, relative to control plants (Figure 6B). Relative to leaves inoculated with BSMV:γ, the transcription of the endogenous *TaCaM3* was successfully silenced in the fourth leaves of BSMV:TaCaM3 plants (Figure 6C). Moreover, we microscopically examined *TaCaM3*-silenced leaves infected with *Pst* CYR23. The length of the hyphae and the numbers of hyphal branches, haustorial mother cells, and haustoria at 24 hpi were significantly greater in the *TaCaM3*-silenced plants than the control plants (Figures 6D and E). At 120 hpi, the infection areas of *Pst* in *TaCaM3*-silenced plants were much larger than those in the control plants (Figure 6F). In addition, to study the effects of the knockdown of *TaCaM3* on the expression of defense-related genes in wheat seedling leaves inoculated with CYR23, we assayed the expression of three PR genes in *TaCaM3*-silenced plants. The transcriptions of *TaPR1*, *TaPR2*, and *TaPR5* were reduced at 24 and 48 hpi in the *TaCaM3*-knockdown plants (Figure 6G), indicating that silencing of *TaCaM3* may reduce the resistance of wheat.

**Figure 6.**
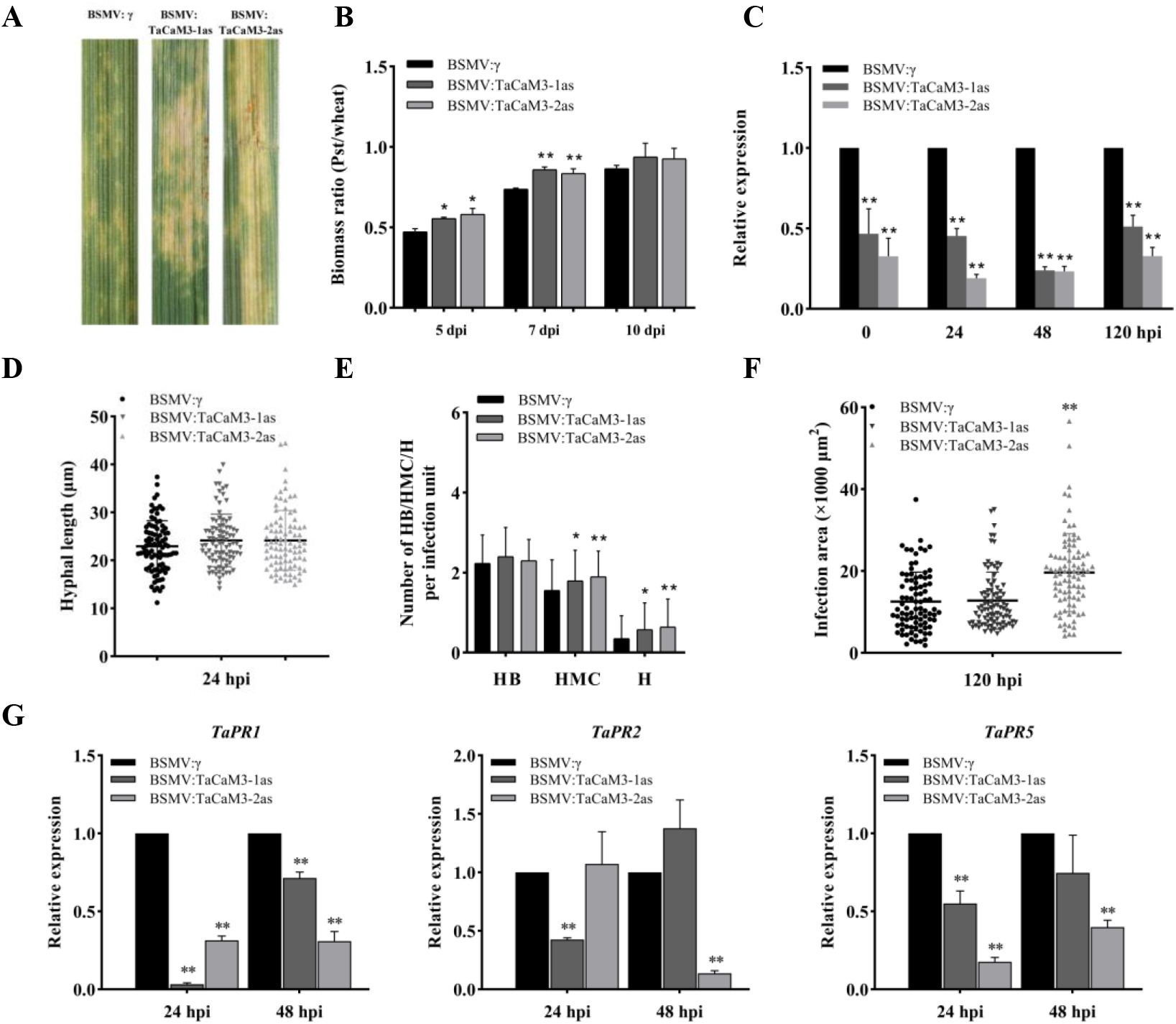
Silencing *TaCaM3* reduces wheat resistance to *Pst* CYR23. A, Disease phenotypes of the fourth leaves pre-inoculated with BSMV constructs and then challenged with *Pst* CYR23. B, The biomass ratio (*Pst*/wheat) measurements of total DNA extracted from BSMV-treated wheat leaves infected by CYR23 at 5, 7 and 10 dpi based on qRT-PCR. Ratio of total fungal DNA to total wheat DNA was assessed by normalizing the data to the wheat gene *TaEF-1α* and the *Pst* gene *PstEF1*. C, The relative expression levels of *TaCaM3* in leaves inoculated with CYR23 were assayed by qRT-PCR at 0, 24, 48, and 120 hpi. D, The hyphal length of *Pst* in *TaCaM3*-silenced plants at 24 hpi. E, Average number of hyphal branches (HB), haustorial mother cells (HMC), and haustoria (H) of *Pst* in each infection site were counted at 24 hpi. F, *TaCaM3*-silenced plants show a significant increase in infection unit area at 120 hpi. Means were calculated from 50 infection sites of three biological replicates and were represented as solid lines in the picture. G, Relative expression of the marked defense-related genes in the fourth leaves at 24 and 48 hpi. Transcript levels were quantified by qRT-PCR and normalized with *TaEF-1α*. The mean and standard deviation were calculated from three independent biological replications. Asterisks indicate a significant difference (*P < 0.05, **P < 0.01) from BSMV:γ inoculated plants using the Student’s *t* test.

Because of the high transcriptional levels of *TaCaM3* in compatible interactions, we also used the CYR31 to inoculate *TaCaM3*-knockdown plants. Although all leaves inoculated with CYR31 produced numerous uredospores (Figure 7A), the fungal biomass was significantly increased in BSMV:TaCaM3 leaves at 5 dpi compared to the control plants (Figure 7B). The silencing efficiency analyses performed by RT-qPCR indicated that *TaCaM3* was significantly knocked down in leaves inoculated with CYR31 at 0, 24, 48, and 120 hpi (Supplemental Figure S9). Moreover, histological observations showed that the hyphal length at 24 hpi (Figure 7C) and infection areas of *Pst* at 120 hpi (Figure 7D) were also greater after the silencing of *TaCaM3* than plants inoculated with BSMV:γ. These results also confirmed that silencing of *TaCaM3* reduced the wheat resistance to *Pst*, and *TaCaM3* may contribute to the basal immune response to *Pst* in wheat.

**Figure 7.**
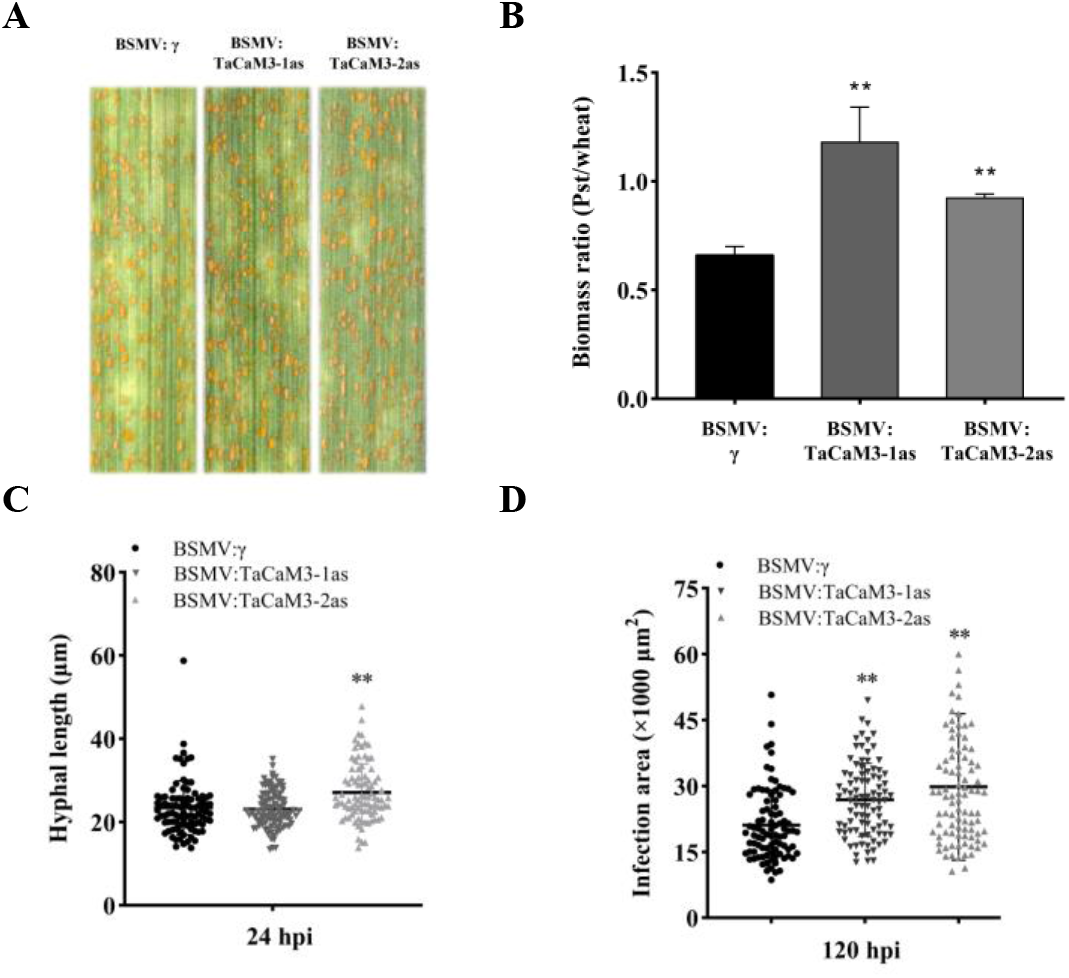
Knocking down *TaCaM3* enhances wheat susceptibility to *Pst* CYR31. A, Disease phenotypes of the fourth leaves pre-inoculated with BSMV constructs and then challenged with *Pst* CYR31. B, The biomass ratio (*Pst*/wheat) measurements of total DNA extracted from BSMV-treated wheat leaves infected by CYR31 at 5 dpi based on qRT-PCR. C, Significant increase in the hyphal length of *Pst* in *TaCaM3*-silenced plants at 24 hpi. E, *TaCaM3*-silenced plants show a significant increase in infection unit area at 120 hpi. Means were calculated from 50 infection sites of three biological replicates and were represented as solid lines in the picture. Asterisks indicate a significant difference (**P < 0.01) from BSMV:γ inoculated plants using the Student’s *t* test.

### TaBln1 impairs Ca^2+^ influx possibly by interacting with TaCaM3

CaMs, as a universal Ca^2+^ sensor, can transmit the initial signal of cytosolic Ca^2+^ elevation (upon pathogen perception) to downstream targets in a signal transduction cascade (Garcia-Brugger et al., 2006). In this study, to further study the function of *TaCaM3*, two PAMPs (chitin and flg22) treatments were selected to measure the *TaCaM3* transcript levels by RT-qPCR. The results of RT-qPCR showed that *TaCaM3* was only upregulated upon chitin treatment (Figure 8A) but did not change upon flg22 treatment (Figure 8B). This indicated that chitin treatment could be used for further study.

**Figure 8.**
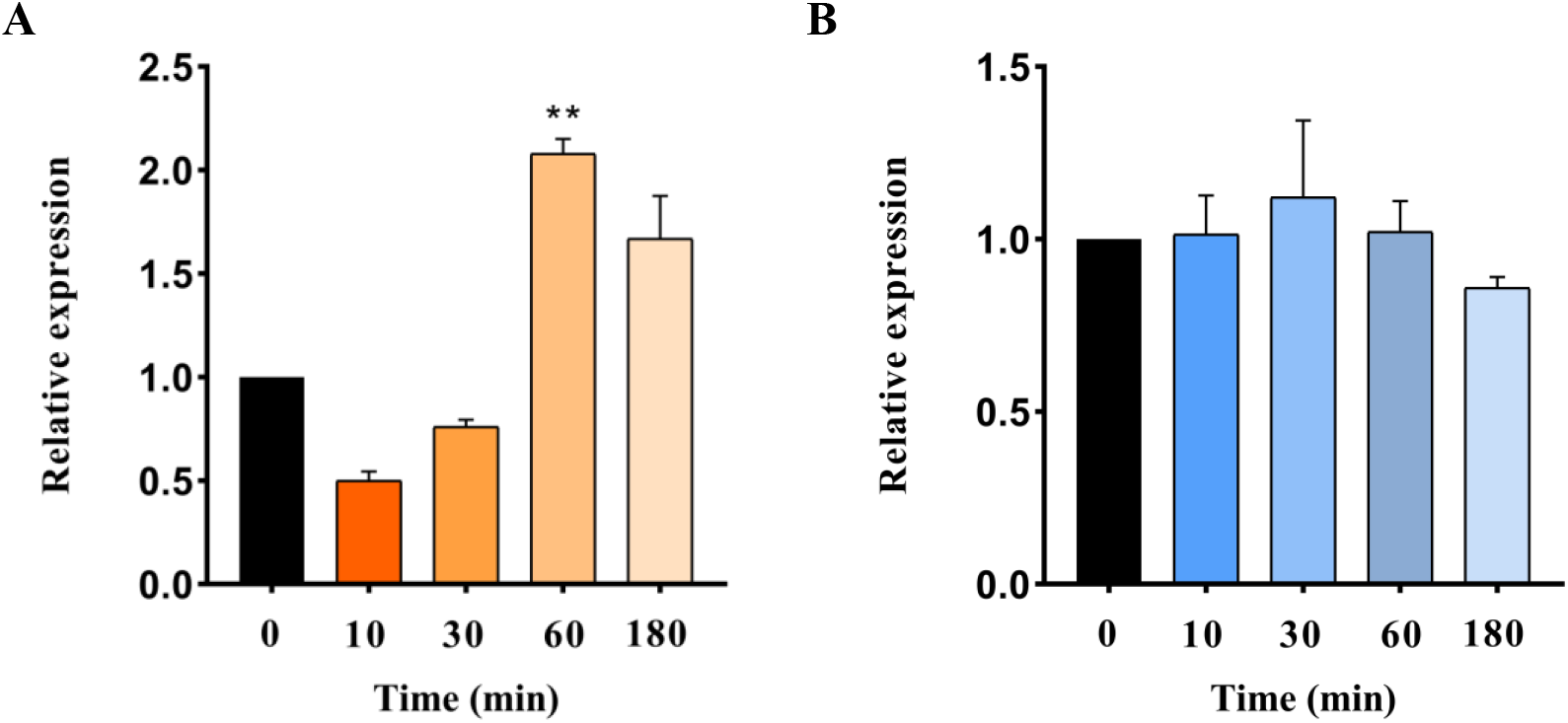
*TaCaM3* expression changes in response to two PAMPs (chitin and flg22) treatment. A, The expression levels of *TaCaM3* in leaves treatment with chitin were assayed by qRT-PCR at 0, 10, 30, 60 and 180 min post treatment. B, The expression levels of *TaCaM3* in leaves treatment with flg22 were assayed by qRT-PCR at 0, 10, 30, 60 and 180 min post treatment. Expression levels were normalized to *TaEF-1α*. The mean and standard deviation were calculated from three independent biological replications. Asterisks indicate a significant difference (**P < 0.01) from control (water treatment) using the Student’s *t* test.

We also investigated whether *TaCaM3* may affect Ca^2+^ influx by measuring the dynamics of Ca^2+^ flux in mesophyll cells after treatment with chitin *in vivo*, using a NMT. All of the leaves tested were clearly responsive to chitin treatment. In response to chitin stimulation, the mesophyll cells of control plants inoculated with BSMV:γ exhibited robust Ca^2+^ influx; however, little change was seen in *TaCaM3* knockdown plants (Figure 9A). Because TaBln1 can interact with TaCaM3, we speculated that TaBln1 might also affect the Ca^2+^ influx by influencing TaCaM3. To test this hypothesis, we also measured the Ca^2+^ influx in *TaBln1* knockdown plants. The results showed that the mesophyll cells of plants inoculated with BSMV:TaBln1, but not those in control plants, exhibited stronger and more rapid Ca^2+^ influx (Figure 9B). These results suggest that TaCaM3 could affect Ca^2+^ influx, and this ability was impaired by interaction with TaBln1.

**Figure 9.**
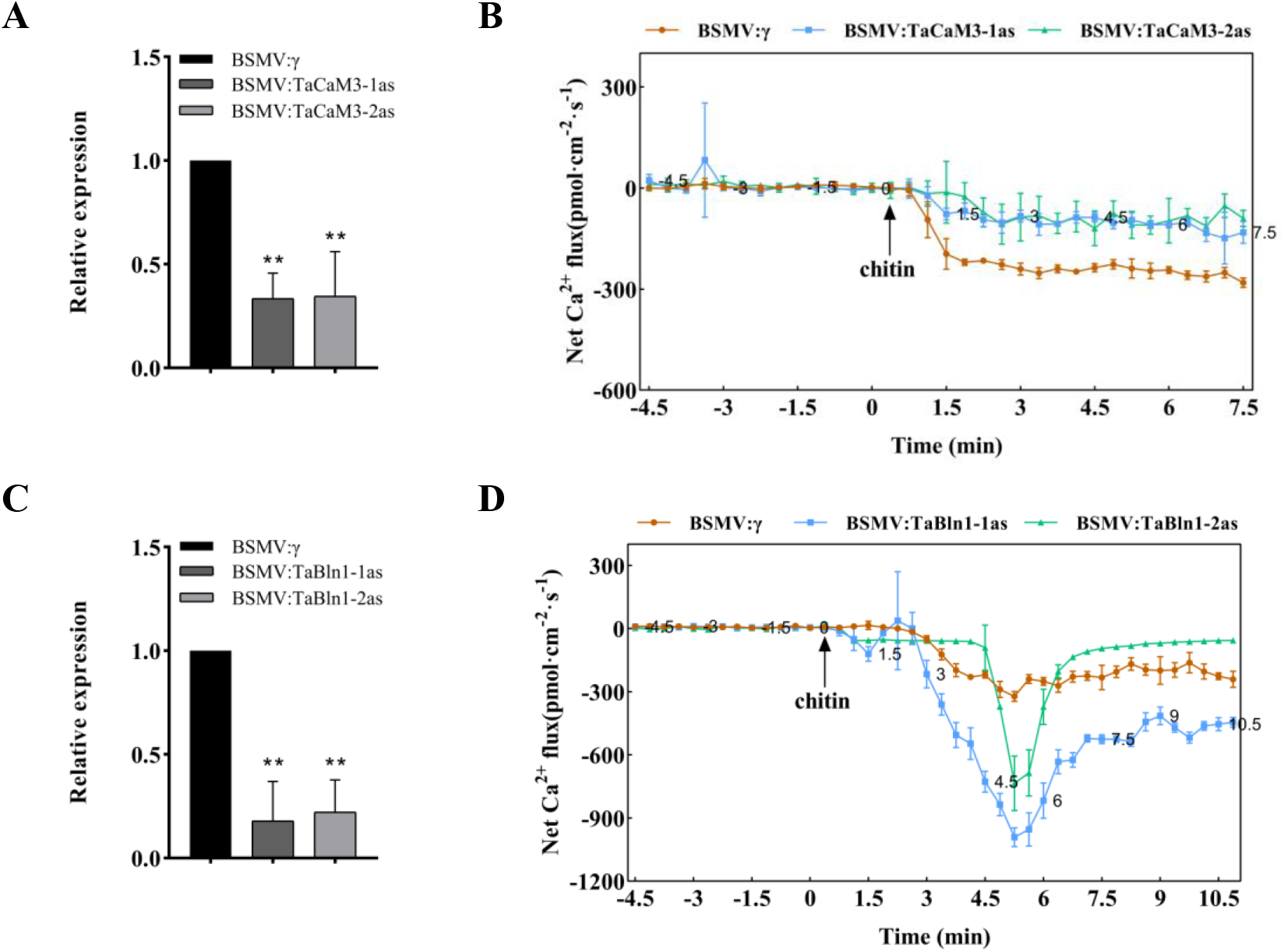
Ca^2+^ signaling in response to chitin in *TaBln1*- and *TaCaM3*-knockdown plants. A, The expression of *TaCaM3* was substantially knocked down by VIGS. B, Comparison of calcium influx in mesophyll cells from BSMV:γ inoculated plants and BSMV:TaCaM3 inoculated plants after chitin treatment. C, The expression of *TaBln1* was substantially knocked down by VIGS. D, Comparison of calcium influx in mesophyll cells from BSMV:γ inoculated plants and BSMV:TaBln1 inoculated plants after chitin treatment. Transcript levels were quantified by qRT-PCR and normalized with TaEF-lα. The relative expression of *TaBln1* and *TaCaM3* were calculated using the comparative threshold method (2^-ΔΔCT^). Values were derived from three biological repetitions. Asterisks indicate a significant difference (**P < 0.01) from BSMV:γ inoculated plants using the Student’s *t* test.

## Discussion

The use of loss-of-function mutations in S genes can save expenditure on pesticides and fungicides and eliminate the negative impacts on the environment that such approaches may have. Hence, the study of S genes is of great interest for engineering plant disease resistance. In this study, we identified *TaBln1* in wheat, which had high expression levels in the early stages of a compatible interaction but showed no significant change in an incompatible interaction. Thus, we speculated that *TaBln1* may play a role in wheat susceptibility. Similar findings regarding *Bln* genes have been reported in barley: the overexpression *Bln1* and *Bln2* by BSMV significantly increased susceptibility of barley to *Bgh* (Meng et al., 2009; Xu et al., 2015), whereas the silencing of *Bln1* enhanced barley resistance in compatible interactions (Meng et al., 2009). In this study, we found that silencing *TaBln1* limited the hyphal growth and fungal colony areas of *Pst* and increased ROS accumulation of wheat, resulting in a reduction in the number of *Pst* uredinia. Thus, we demonstrated that the function of the *TaBln1* gene is similar to the *Bln1* and *Bln2* genes of barley, which played a negative regulatory role in the interaction between wheat and *Pst*. Interestingly, the *Bln* genes may be unique to the cereal grain crops barley, wheat, and rice (Meng et al., 2009). The identification of *TaBln1*, as a negative regulatory factor, would provide a preliminary target in the future for editing *Bln* genes with CRISPR to achieve durable disease control.

Previous studies have indicated that Bln family members in barley are cysteine-rich small peptides (Meng et al., 2009), and both Bln1 and Bln2 interact with CaM (Xu et al., 2015). In this study, we compared the amino acids of Bln1 in wheat with Bln1 and Bln2 in barley and found that they all had conserved cysteines (Supplemental Figure S10). These conserved cysteines in CaM targets are important for CaM–target interactions (Moore et al., 1999). Thus, we speculated that TaBln1 possessed a function somewhat similar to HvBln1 and HvBln2, which may interact physically with CaM. Interestingly, our results of BiFC, pulldown, LUC, and Co-IP analyses are in support of this speculation. This interaction between TaBln1 and TaCaM3 implies that they might be regulated differently during wheat–*Pst* interactions. We speculate that *TaBln1* negatively regulates immunity in wheat by affecting the function of *TaCaM3*.

CaMs play a critical role in plant defense (Takabatake et al., 2007; Zhu et al., 2010). TaCaM3, a CaM, is a highly conserved intracellular calcium sensor (Bouché et al., 2005; Defalco et al., 2009). In this study, we identified *TaCaM3* as a positive regulator of basal immunity against the *Pst* fungus, and we found that transiently silencing *TaCaM3* in wheat decreased resistance to fungal infection, which increased areas of infection and sporulation. Numerous studies have suggested a key role for CaM-mediated calcium signaling in plant growth and stress responses (Liu et al., 2003; Du et al., 2009). Thus, we speculated that *TaCaM3* plays a positive regulatory role in the interaction between wheat and *Pst* through TaCaM3-mediated calcium signaling. To test this hypothesis, we chose chitin from two PAMPs (chitin and flg22) as a great substitute for *Pst*, which has certain limitations, for further study.

Intracellular calcium transients during plant–pathogen interactions are necessary early events that lead to local and systemic acquired resistance (Lecourieux et al., 2006). Calcium binding to CaMs regulates cellular processes directly by binding to specific DNA sequences and modulates gene expression indirectly by interacting with other proteins in a Ca^2+^-dependent manner and modulating their activity (Reddy et al., 2011). Therefore, intracellular calcium transients are essential for CaMs to function in plant defense signaling pathways. In this study, we measured the dynamics of Ca^2+^ flux in mesophyll cells after treatment with chitin *in vivo* using NMT. The results indicated that transiently silencing *TaCaM3* decreased the Ca^2+^ influx induced by chitin, consistent with the results of VIGS experiments. We speculated that transient silencing of *TaCaM3* affected the balance between calcium binding and calcium influx and ultimately weakened the disease resistance of wheat plants. Previous studies have indicated that CaM induces a net Ca^2+^ influx into protoplasts, leading to an increase in cytoplasmic Ca^2+^ levels (Wang et al., 2009a). In line with this finding, our results indicated that TaCaM3 affected Ca^2+^ influx. In addition, we proved that TaBln1 could interact with TaCaM3, so we speculated that TaBln1 might also affect Ca^2+^ influx by affecting the function of TaCaM3. Interestingly, transiently silencing *TaBln1* produced a stronger and more rapid Ca^2+^ influx, which was also consistent with the results of VIGS. Recently, similar findings on the relationship between Ca^2+^ influx and immune responses have been reported. For example, by comparing the calcium influx in the mesophyll cells of WT and transgenic rice plants after chitin or flg22 treatments, a study found that activated OsCNGC9 induced extracellular Ca^2+^ influx and then triggered ROS burst and PTI-related gene expressions, which ultimately led to enhanced disease resistance in rice (Wang et al., 2019). Therefore, we speculated that TaCaM3 could affect Ca^2+^ influx, and this ability was impaired by the interaction with TaBln1. TaCaM3 could not effectively transmit the initial signal of cytosolic Ca^2+^ elevation to downstream targets in a signal transduction cascade, which led to a weakening of plant disease resistance. To confirm our speculation, the relative expression of *TaCaM3* in *TaBln1*-silenced leaves was determined by RT-qPCR. As expected, silencing of *TaBln1* increased the transcription of *TaCaM3* (Supplemental Figure S11).

In addition, analyses of the *TaBln1* sequence found that *TaBln1* is a three-copy gene located on chromosomes 4A, 5B, and 5D. Because T (4AL; 5AL) exists in diploid *T. monococcum* (Dubcovsky et al., 1996), the evolution of the wild emmer wheat chromosome 4A must have started with this translocation. In the evolution of wheat chromosomes 4A, 5A, and 7A, wheat chromosome 4A was subjected to chromosome rearrangements a reciprocal translocation T (4AL; 5AL) (Dvorak et al., 2018). Analyses of the TaBln1 protein structure found that although TaBln1 contains a signal peptide (1-19AA), it also has a putative transmembrane domain (Supplemental Figure S4). In *Arabidopsis*, receptor kinase FLS2 contains a signal peptide and a transmembrane domain, which can be targeted to the plasma membrane (Gómez-Gómez et al., 2000). OsCERK1, which encodes a receptor-like kinase containing a signal peptide, an extracellular domain, a transmembrane region, and an intracellular Ser/Thr kinase domain, is also localized to the plasma membrane (Shimizu et al., 2010). Therefore, we speculate that TaBln1 possess a similar localization to the aforementioned proteins. As expected, this study confirmed the localization of TaBln1 on the plasma membrane and nuclear membrane through tobacco subcellular localization. At the same time, a plasmolysis experiment confirmed the localization of TaBln1 (Figure 5B; Supplemental Figure S12). This result is different from the localization of BLN1 and BLN2, which can be secreted to the apoplast in barley (Xu et al., 2015). Consistent with the negative control free GFP but in contrast with the positive control SP(TaPR1a)-GFP (Bi et al., 2020), TaBln1, after plasmolysis, was still localized to the membrane, and no apoplastic signals were observed (Supplemental Figure S12). We speculated that this may be related to its negative regulatory function in the interaction between wheat and *Pst*. Interestingly, the results of co-localization showed that TaCaM3 and TaBln1 were co-localized to the plasma membrane, and the BIFC results showed that the fluorescence signal is mainly distributed at the plasma membrane. Therefore, we speculate that the interaction between TaBln1 and TaCaM3 mainly occurs at the plasma membrane. TaBln1 affects the calcium influx modulated by TaCaM3, possibly by hijacking TaCaM3 to the plasma membrane.

## Conclusions

The present study reveals the potential molecular mechanism of *TaBln1* in the wheat–*Pst* interaction. In uninoculated wheat leaves (Figure 10A), the expression of *TaBln1* is maintained at a low level. TaCaM3 functions normally in binding Ca^2+^, and it may be transferred to downstream targets to activate the immune responses. During *Pst* infection of wheat (Figure 10B), the expression of *TaBln1* may be upregulated and localized to the plasma membrane. The interaction between TaBln1 and TaCaM3 results in the accumulation of TaCaM3 on the plasma membrane. This, in turn, affects the balance between calcium binding and calcium influx, thereby reducing the transmission of Ca^2+^ signals to downstream targets, and ultimately weakening the disease resistance of wheat. Obviously, a better understanding of the physiological significance of the cytosolic Ca^2+^ changes in plant–pathogen interactions will involve the identification and functional analysis of downstream targets of cytosolic Ca^2+^. Thus, most intracellular target proteins that sense and relay Ca^2+^ signatures toward the appropriate defense responses remain possible objects of future testing.

**Figure 10.**
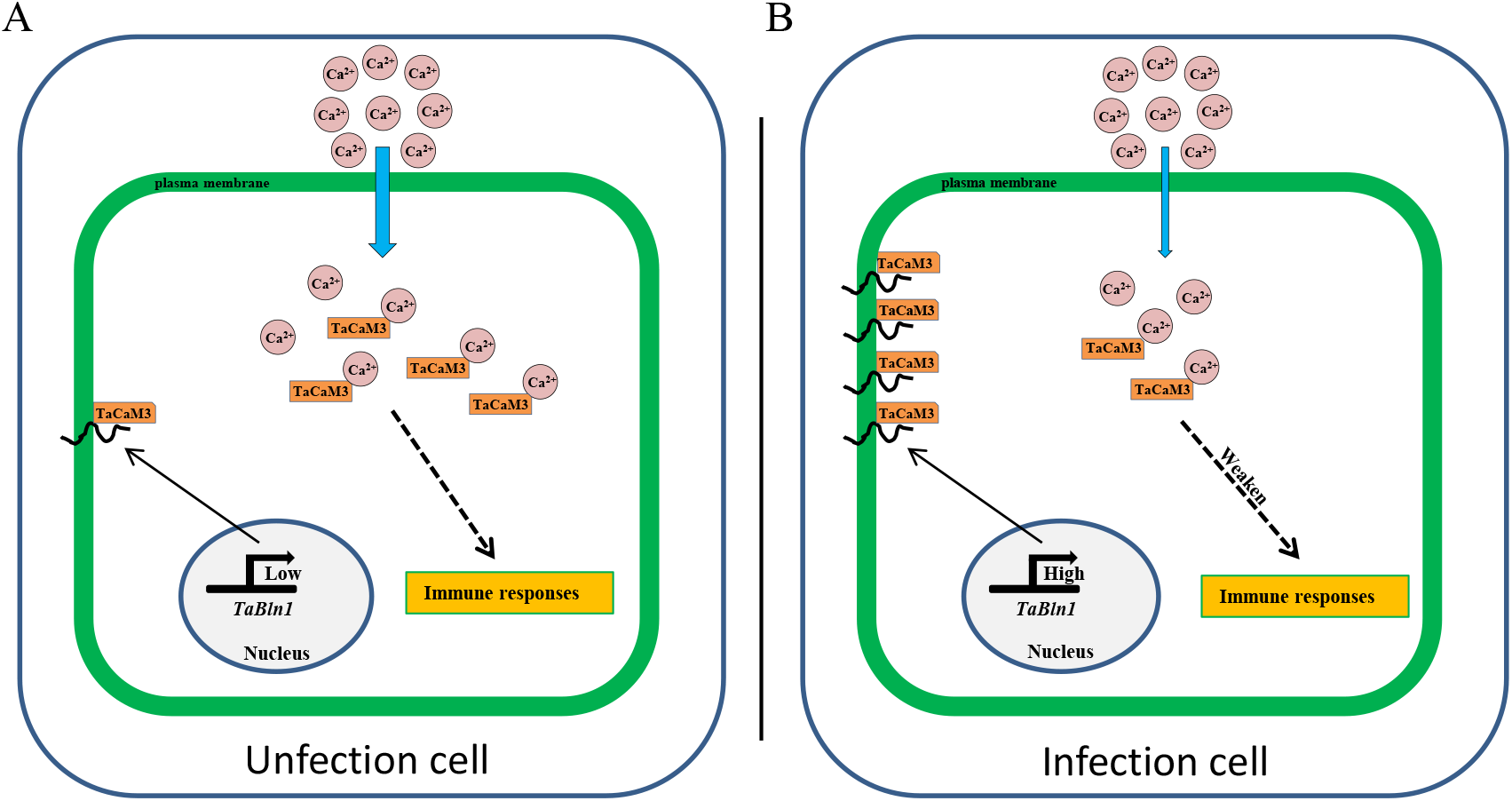
The proposed model of the function of *TaBln1* in the wheat-*Pst* interaction. A, In uninoculated wheat leaves (Figure 10A), the expression of *TaBln1* is maintained at a low level. TaCaM3 functions normally in binding Ca^2+^, and it may be transferred to downstream targets to activate the immune responses. During *Pst* infection of wheat (Figure 10B), the expression of *TaBln1* may be upregulated and localized to the plasma membrane. The interaction between TaBln1 and TaCaM3 results in the accumulation of TaCaM3 on the plasma membrane. This, in turn, affects the balance between calcium binding and calcium influx, thereby reducing the transmission of Ca^2+^ signals to downstream targets, and ultimately weakening the disease resistance of wheat.

## Materials and Methods

### Plant materials, inoculation, and treatment

Wheat (*Triticum aestivum* cv. Suwon 11), *Nicotiana benthamiana*, and two *Pst* pathotypes, CYR23 (avirulent) and CYR31 (virulent), were used in this study. Suwon 11 exhibits a high resistance to CYR23 but high susceptibility to CYR31 (Cao et al., 2003). Wheat seedling cultivation and inoculation with *Pst* were performed as described previously (Kang et al., 2002). Leaves were collected at 0, 12, 24, 48, 72, and 120 post-inoculation (hpi) for RNA isolation (Wang et al., 2007). The *N. benthamiana* was grown at 22°C and used for transient expression.

### Genomic DNA, total RNA extraction, and cDNA synthesis

Genomic DNA was extracted using the cetyltrimethylammonium bromide (CTAB) method (Porebski et al., 1997). The total RNA from the wheat leaves was isolated using a Quick RNA isolation Kit (TIANGEN BIOTECH), according to the manufacturer’s instructions. The extracted RNA was reversely transcribed into cDNA using the RevertAid First Strand cDNA Synthesis Kit (Thermo Scientific) and oligo (dT)_18_ primer following the manufacturer’s protocol.

### Cloning of *TaBln1* and *TaCam3* and sequence analysis

To clone the *TaBln1* and *TaCam3* sequences, specific primers (Supplementary Table S1) were designed using the Primer 5.0 software. The *TaBln1* and *TaCam3* sequences were PCR-amplified using Suwon 11 cDNA sample as a template. The obtained sequences were aligned using the *T. aestivum* cv. Chinese Spring (CS) genome from the Ensemble Plants database (http://plants.ensembl.org/Multi/Tools/Blast) and the NCBI (http://www.ncbi.nlm.nih.gov/). The multi-sequence alignments were carried out using DNAMAN 8 software. The molecular sizes of *TaBln1 and TaCam3* were predicted using the Compute pI/Mw tool (https://web.expasy.org/compute_pi/). The signal peptide of TaBln1 was identified using SignalP 5.0 (http://www.cbs.dtu.dk/services/SignalP/). The transmembrane domain of TaBln1 was identified using TMHMM Server v. 2.0 (http://www.cbs.dtu.dk/services/TMHMM/).

### Plasmid construction

To construct the vector of subcellular localization, the coding sequences for *TaBln1* and *TaCaM3* without a stop codon were amplified and inserted into the *Nco*I and *Spe*I restriction sites of the plant binary expression vector pCAMBIA1302 containing the GFP-tag sequence, respectively. In addition, the *TaBln1* without a stop codon was inserted into the pICH86988 vector with the mCherry tag.

For plasmid constructs for the silencing system, two small cDNA fragments based on the results of a BLASTN search of the NCBI (http://www.ncbi.nlm.nih.gov/) that showed the lowest sequence similarity with other wheat genes were inserted into *Pac*I and *Not*I restriction sites of the virus plasmid γ, respectively (Holzberg et al., 2002).

For the interaction analyses of TaBln1 and TaCam3 using BIFC, *TaBln1* and *TaCam3* were subcloned into pSPYNE(R)173 and pSPYCE(M) with *Bam*HI and *Xho*I restriction sites to generate the pSPYNE(R)173-*TaBln1* and pSPYCE(M)-*TaCam3* vectors, respectively (Waadt et al., 2008). The coding sequence of *TaBln1* was inserted into the *Spe*I restriction site of pBinGFP2 vector and *TaCam3* with HA tag was inserted into the *Bsa*I restriction site of pICH86988 vector to generate pBinGFP2-*TaBln1* and pICH86988-*TaCam3* for co-immunoprecipitation. The coding sequences of *TaBln1* and *TaCam3* were subcloned into the N- or C-terminal fragment of LUC (nLUC or cLUC) with *Bam*HI and *Sal*I restriction sites to generate the *TaBln1*-nLUC and *TaCam3*-cLUC vectors for split-luciferase assays, respectively. The coding sequence of *TaBln1* was subcloned into the *Eco*RI and *Xho*I restriction sites of the pGEX-4T-1 expression vector containing the GST-tag sequence; the coding sequence of *TaCam3* was subcloned into the *Eco*RI and *Xho*I restriction sites of the pET-28a vectors containing the His-tag sequence for the pulldown assay. The primers for all of the plasmid constructions are listed in Supplemental Table S1.

### BSMV-mediated gene silencing

To transiently silence *TaBln1* and *TaCam3*, the corresponding vectors (*TaPDS*-γ, recombinant γ-gene and γ), α and β were linearized using corresponding enzymes and transcribed into RNA. The transcripts of α and β were mixed in a 1:1:1 ratio with transcripts of *TaPDS*-γ, recombinant γ-gene, and γ in FES buffer (0.1 M glycine pH 8.9, 0.06 M K_2_HPO_4_, 1% bentonite, 1% sodium pyrophosphate and 1% celite), respectively. Then the second wheat leaves were inoculated with the BSMV RNA mixture and maintained at 26°C (Holzberg et al., 2002). BSMV: TaPDS (wheat phytoene desaturase) was used as a positive control. When the virus phenotype (photobleaching in positive control) was observed (12 days after BSMV treatment), the fourth leaves were inoculated with *Pst* CYR23 or CYR31 and samples were harvested at 0, 24, 48, and 120 hpi for histological observation and RNA isolation. RNA was used to confirm the silencing efficiency for each assay by RT-qPCR. The phenotypes were recorded, and representative photographs were captured at 14 dpi. Three independent sets of inoculations were performed, consisting of 60 seedlings inoculated for each BSMV virus.

### Histological analyses of silenced plants inoculated with *Pst*

For the histological observations of fungal growth, the samples of silenced plants were cut into several fragments and decolorized in destaining solution (absolute ethyl alcohol:acetic acid, 1:1 v/v) as previously described (Cheng et al., 2015). Then these fragments were decolorized in chloral hydrate for 24 h followed with autoclaving at 121°C for 2 min. Wheat germ agglutinin (WGA) conjugated to Alexa Fluor 488 (Invitrogen) was used to stain the *Pst* infection structures as previously described (Ayliffe et al., 2011).

For the detection of H_2_O_2_ accumulation, infected leaves were cut and the ends were immersed in a solution containing 1 mg/mL 3,3-diaminobenzidine (DAB, Amresco) for 4–6 h (Wang et al., 2007). Then these fragments were decolorized in destaining solution to remove the chlorophyll. Only those infected sites where an appressorium had formed over a stoma were considered a successful penetration site of *Pst*. The accumulation of H_2_O_2_, hyphal length, infection area, and a number of hyphal branches, haustorial mother cells, and haustoria were observed with an Olympus BX-51 microscope (Olympus) and measured using the cellSens Entry software (Olympus).

### Protein interaction assays

For the BiFC assay, the pSPYNE(R)173-TaBln1 and pSPYCE(M)-TaCam3 vectors were transformed into *A. tumefaciens* GV3101. The *Agrobacterium* strains were co-infiltrated at an OD_600_ of 0.6 into *N. benthamiana* leaves. Two days after inoculation, YFP fluorescence was captured by confocal microscopy with a 488 nm laser.

For the Co-IP assay, the pBinGFP2-TaBln1 and pICH86988-TaCam3 vectors were transformed into *A. tumefaciens* GV3101 and co-infiltrated into *N. benthamiana* leaves. The pBinGFP2 empty vector was used as the negative control. Two days after inoculation, the infiltrated leaves were harvested and ground to a powder in liquid nitrogen and then homogenized in RIPA buffer (50 mM Tris-HCl, pH 8, 150 mM NaCl, 1% NP-40, 0.25% sodium deoxycholate, and 0.1% SDS) with 1 mM PMSF (Beyotime Biotechnology). Then the extract was centrifuged at 12000 rpm for 10 min, and the supernatant was transferred into a fresh tube for the Co-IP assay. For immunodetection, the supernatant was mixed with 15 μL GFP-trap A beads (Chromotek) and incubated at 4°C for 1.5 h. The beads were collected and washed five times with 500 μL wash buffer (50 mM Tris-HCl, pH 7.4, 150 mM NaCl and 0.5% Tween-20). Proteins bound to the beads were boiled for 10 min and detected by Western blotting with anti-HA and anti-GFP, respectively. The monoclonal antibodies anti-HA (Beyotime Biotechnology) and anti-GFP (Beyotime Biotechnology) were used at a 1:5,000 dilution and followed by incubation with a second antibody, anti-mouse Ig-horseradish peroxidase (1:2000; Beyotime Biotechnology). Protein bands were detected using SuperSignal West Femto Maximum Sensitivity substrate (Thermo Scientific).

Split-luciferase assays were performed as previously described (Chen et al., 2008). *A. tumefaciens* GV3101 with different constructs were co-infiltrated into *N. benthamiana* leaves, respectively. After 2 days, 1 mM luciferin (AbMole) was sprayed onto the inoculated leaves, and the LUC activity was captured with a PlantView100 assay system (BLT PHOTON TECHNOLOGY).

For the pulldown assay, GST fusion proteins of TaBln1 and His fusion proteins of TaCam3 were expressed in *E. coli* (BL21) by induction with 0.3 mM IPTG at 16°C overnight. Crude proteins of TaBln1 with GST tag were purified using standard techniques with glutathione-sepharose following a previous study (Harper et al., 2001), and TaCam3 with the His tag was purified via Ni-chelating affinity chromatography. Equal amounts of GST-tagged TaBln1 or GST (negative control) were mixed with TaCam3-His and incubated at 4°C for 2 h with GST beads (Thermo Scientific). The beads were collected and washed five times using the GST Protein Interaction Pull down Kit (Thermo Scientific) according to the manufacturer’s instructions to detect the recovered TaCam3-His levels. The proteins bound to the beads were boiled for 10 min and detected by Western blotting with anti-GST and anti-His antibodies. Monoclonal anti-GST (Beyotime Biotechnology) and anti-His (Beyotime Biotechnology) antibodies were used at a 1:5,000 dilution and followed by incubation with a second antibody, anti-mouse Ig-horseradish peroxidase (Beyotime Biotechnology). The protein bands were detected using SuperSignal West Femto Maximum Sensitivity substrate (Thermo Scientific).

### Subcellular localization of TaBln1 and TaCam3 in *N. benthamiana*

To determine the subcellular localization of TaBln1 and TaCam3, *Agrobacterium* strains GV3101 harboring TaBln1-GFP and TaWpi6-mCherry, a positive control marker protein that is localized to the plasma membrane (Imai et al., 2005), were co-infiltrated into 4-week-old *N. benthamiana* leaves. To determine the co-localization of TaBln1 and TaCam3, *N. benthamiana* leaves were infiltrated with GV3101 carrying TaBln1-mCherry and TaCam3-GFP. The infiltrated seedlings were transferred to a growth chamber at 22°C with a 16 h light photoperiod. GFP fluorescence signals, with an excitation laser at 488 nm and mCherry fluorescence signals with an excitation laser at 546 nm were monitored at 48 h post transformation using an Olympus FV3000 Confocal Laser Microscope. All of the assays were performed in duplicate and repeated at least three times.

### RT-qPCR analyses

Transcription levels were analyzed using a Real-Time PCR Detection System (Bio-Rad). Primer designs and RT-qPCR reactions were conducted as described previously (Wang et al., 2009b). The wheat elongation factor *TaEF-1a* gene was used as a wheat internal reference. Fungal biomass changes in the *Pst*-infected wheat leaves were analyzed as described previously (Chang et al., 2017). Biomass was determined by absolute quantification using a double-standard curve (Lee et al., 2008). The constitutively expressed wheat stripe rust elongation factor gene *PstEF1* was used for the quantification of wheat stripe rust fungus (Yin et al., 2009). All of the primers used are listed in Supplemental Table S1. RT-qPCR analyses used data from three biological repeats, with each group containing three technical repeats. Relative expression was estimated using the 2^−ΔΔCT^ method (Livak et al., 2001).

### Quantification of PAMP-induced gene expression and measurements of net Ca^2+^ flux

The detection of PAMP-induced gene expression was conducted as previously described, with minor modifications (Park et al., 2012; Schoonbeek et al., 2015). Four 2 cm long strips of wheat leaves were cut and placed into 2 mL tubes with water and preinfiltrated by vacuum for 3× 1 min. Then the materials were left to recover in a growth cabinet overnight to recover from wounding stress and water infiltration. Afterward, water was replaced by fresh water (mock) or 10 μM flg22 peptide (Bankpeptide) or 10 μM chitin (Santa Cruz. Biotechnology) for 10, 30, 60, or 180 min, followed by freezing in liquid nitrogen. Materials were used for RT-qPCR assays with the primers listed in Supplemental Table S1.

Net Ca^2+^ flux was measured using the NMT-YG-100 (YoungerUSA) as previously described (Ma et al., 2015; Wang et al., 2019). In brief, leaves sampled from transiently silenced seedling were immobilized in measuring buffer (0.1 mM KCl, 0.1 mM CaCl_2_, 0.1 mM MgCl_2_, 0.5 mM NaCl, 0.3 mM MES, 0.2 mM Na_2_SO_4_, and pH 6.0) for 30 min equilibration. The steady-state fluxes in leaf mesophyll cells were continuously recorded for 5 min prior to the chitin treatments. Thereafter, the chitin was slowly added to the measuring buffer until the chitin concentration reached 10 μM. Then the transient flux of Ca^2+^ was recorded and continued for 10 min.

### Data analyses

The data were analyzed using Student’s *t*-test, with GraphPad Prism 8.0 statistical software to determine the significant differences between control and treatment.

### Accession numbers

Sequence data from this article can be found in the National Center for Biotechnology database (http://www.ncbi.nlm.nih.gov/) and the Ensembl Plants portal (http://plants.ensembl.org/Triticum_aestivum/Info/Index) under the following accession numbers: *TaBln1-4A* (AK333112.1), *TaBln1-5B* (xxx), *TaBln1-5D* (xxx), *TaCaM3-2A* (TraesCS2A02G098100), *TaCaM3-2B* (TraesCS2B02G113800), *TaCaM3-2D* (TraesCS2D02G097500), *TaCaM3-4A* (TraesCS4A02G126700), *TaCaM3-4B* (TraesCS4B02G178200), *TaCaM3-4D* (TraesCS4D02G179800), *TaEF-1α* (Q03033). *TaPR1* (AF384143), *TaPR2* (DQ090946), *TaPR5*, (FG618781).

## List of authors’ contributions

S.Y.G., Z.S.K., X.J.W., and X.M.Z. conceived and designed the experiments; Q.Z., Q.L.X., T.L. contributed to the original concept of the project. S.Y.G., Y.Q.Z., P.Z., M.L., X.L. performed the experiments; S.Y.G. and X.L. performed the data analyses. S.Y.G. wrote the manuscript. Z.S.K., X.J.W., and X.M.Z. revised the manuscript.

## References

Acevedo-Garcia J, Kusch S, Panstruga R (2015) Magical mystery tour: MLO proteins in plant immunity and beyond. New Phytol 204: 273–281

Ayliffe M, Devilla R, Mago R, White R, Talbot M, Pryor A, Leung H (2011) Nonhost resistance of rice to rust pathogens. Mol Plant Microbe Interact 24: 1143–1155

Bi W, Zhao S, Zhao J, Su J, Yu X, Liu D, Kang Z, Wang X, Wang X (2020) Rust effector PNPi interacting with wheat TaPR1a attenuates plant defense response. Phytopathology Res 2: 1–14

Bouché N, Yellin A, Snedden WA, Fromm H (2005) Plant-specific calmodulin-binding proteins. Annu Rev Plant Biol 56: 435–466

Caldo RA, Nettleton D, Wise RP (2004) Interaction-dependent gene expression in Mla-specified response to barley powdery mildew. Plant Cell 16: 2514–2528

Cao Z, Jing J, Wang M, Shang H, Li Z (2003) Relation analysis of stripe rust resistance gene in wheat important cultivar suwon 11,suwon 92 and hybrid 46. Acta Botanica Boreali-Occidentalia Sinica 23: 64–68

Chang Q, Liu J, Lin X, Hu S, Yang Y, Li D, Chen L, Huai B, Huang L, Voegele RT, et al. (2017) A unique invertase is important for sugar absorption of an obligate biotrophic pathogen during infection. New Phytol 215: 1548–1561

Chen H, Zou Y, Shang Y, Lin H, Wang Y, Cai R, Tang X, Zhou J-M (2008) Firefly luciferase complementation imaging assay for protein-protein interactions in plants. Plant Physiol 146: 368–376

Cheng Y, Wang X, Yao J, Voegele RT, Zhang Y, Wang W, Huang L, Kang Z (2015) Characterization of protein kinase PsSRPKL, a novel pathogenicity factor in the wheat stripe rust fungus. Environ Microbiol 17: 2601–2617

Dangl JL, Dietrich RA, Richberg MH (1996) Death don’t have no mercy cell death programs in plant-microbe interactions. Plant Cell 8: 1793–1807

Defalco TA, Bender KW, Snedden WA (2009) Breaking the code: Ca^2+^ sensors in plant signalling. Biochem J 425: 27–40

Du L, Ali GS, Simons KA, Hou J, Yang T, Reddy ASN, Poovaiah BW (2009) Ca^2+^/calmodulin regulates salicylic-acid-mediated plant immunity. Nature 457: 1154–1158

Dvorak J, Wang L, Zhu T, Jorgensen CM, Luo MC, Deal KR, Gu YQ, Gill BS, Distelfeld A, Devos KM, et al. (2018) Reassessment of the evolution of wheat chromosomes 4A, 5A, and 7B. Theor Appl Genet 131: 2451–2462

Ellis JG, Lagudah ES, Spielmeyer W, Dodds PN (2014) The past, present and future of breeding rust resistant wheat. Front Plant Sci 5: 1–13

Fukuoka S, Saka N, Koga H, Ono K, Shimizu T, Ebana K, Hayashi N, Takahashi A, Hirochika H, Okuno K, et al. (2009) Loss of function of a proline-containing protein confers durable disease resistance in rice. Science 325: 998–1001

Garcia-Brugger A, Lamotte O, Vandelle E, Bourque S, Lecourieux D, Poinssot B, Wendehenne D, Pugin A (2006) Early signalingevents induced by elicitors of plant defenses. Mol Plant Microbe Interact 19: 711–724

Gómez-Gómez L, Boller T (2000) FLS2: an LRR receptor-like kinase involved in the perception of the bacterial elicitor flagellin in Arabidopsis. Mol Cell 5: 1003–1011

Harper S, Speicher DW (2001) Expression and purification of gst fusion proteins. Curr Protoc Protein Sci Chapter 6: Unit 6.6

Heo WD, Lee SH, Kim MC, Kim JC, Chung WS, Chun HJ, Lee KJ, Park CY, Park HC, Choi JY, et al. (1999) Involvement of specific calmodulin isoforms in salicylic acidindependent activation of plant disease resistance responses. Proc Natl Acad Sci USA 96: 766–771

Hilleary R, Paez-Valencia J, Vens C, Toyota M, Palmgren M, Gilroy S (2020) Tonoplast-localized Ca^2+^ pumps regulate Ca^2+^ signals during pattern-triggered immunity in Arabidopsis thaliana. Proc Natl Acad Sci USA 117: 18849–18857

Holzberg S, Brosio P, Gross C, Pogue GP (2002) Barley stripe mosaic virus-induced gene silencing in a monocot plant. Plant J 30: 315–327

Imai R, Koike M, Sutoh K, Kawakami A, Torada A, Oono K (2005) Molecular characterization of a cold-induced plasma membrane protein gene from wheat. Mol Genet Genomics 274: 445–453

Jørgensen JH (1992) Discovery, characterization and exploitation of Mlo powdery mildew resistance in barley. Euphytica 63: 152

Kang Z, Huang L, Buchenauer H (2002) Ultrastructural changes and localization of lignin and callose in compatible and incompatible interactions between wheat and *Puccinia striiformis*. J Plant Dis Protect 109: 25–37

Lapin D, Ackerveken GVD (2013) Susceptibility to plant disease: more than a failure of host immunity. Trends Plant Sci 18: 546–554

Lecourieux D, Ranjeva R, Pugin A (2006) Calcium in plant defence-signalling pathways. New Phytol 171: 249–269

Lee C, Lee S, Shin SG, Hwang S (2008) Real-time PCR determination of rRNA gene copy number: absolute and relative quantification assays with *Escherichia coli*. Appl Microbiol Biotechnol 78: 371–376

Liu C, Pedersen C, Schultz-Larsen T, Aguilar GB, Madriz-Ordenana K, Hovmøller MS, Thordal-Christensen H (2016) The stripe rust fungal effector PEC6 suppresses pattern-triggered immunity in a host species-independent manner and interacts with adenosine kinases. New Phytol

Liu H, Li B, Shang Z, Li X, Mu R, Sun D, Zhou R (2003) Calmodulin is involved in heat shock signal transduction in wheat. Plant Physiol 132: 1186–1195

Livak KJ, Schmittgen TD (2001) Analysis of relative gene expression data using real-time quantitative PCR and the 2^−ΔΔCT^ Method. Methods 25: 402–408

Ma W, Berkowitz GA (2007) The grateful dead: calcium and cell death in plant innate immunity. Cell Microbiol 9: 2571–2585

Ma W, Berkowitz GA (2011) Ca^2+^ conduction by plant cyclic nucleotide gated channels and associated signaling components in pathogen defense signal transduction cascades. New Phytol 190: 566–572

Ma Y, Dai X, Xu Y, Luo W, Zheng X, Zeng D, Pan Y, Lin X, Liu H, Zhang D, et al. (2015) *COLD1* confers chilling tolerance in rice. Cell 160: 1209–1221

Mccormack E, Braam J (2003) Calmodulins and related potential calcium sensors of Arabidopsis. New Phytol 159: 585–598

Meng Y, Moscou MJ, Wise RP (2009) *Blufensin1* negatively impacts basal defense in response to barley powdery mildew. Plant Physiol 149: 271–285

Moore CP, Zhang J, Hamilton SL (1999) A role for cysteine 3635 of RYR1 in redox modulation and calmodulin binding. J Biol Chem 274: 36831–36834

Nekrasov V, Wang C, Win J, Lanz C, Weigel D, Kamoun S (2017) Rapid generation of a transgene-free powdery mildew resistant tomato by genome deletion. Sci Rep 7: 482

Oliva R, Ji C, Atienza-Grande G, Huguet-Tapia JC, Perez-Quintero A, Li T, Eom J-S, Li C, Nguyen H, Liu B, et al. (2019) Broad-spectrum resistance to bacterial blight in rice using genome editing. Nat Biotechnol 37: 1344–1350

Park CH, Chen S, Shirsekar G, Zhou B, Khang CH, Songkumarn P, Afzal AJ, Ning Y, Wang R, Bellizzi M, et al. (2012) The *Magnaporthe oryzae* effector AvrPiz-t targets the RING E3 ubiquitin ligase APIP6 to suppress pathogen-associated molecular pattern-triggered immunity in rice. Plant Cell 24: 4748–4762

Pavan S, Jacobsen E, Visser RG, Bai Y (2010) Loss of susceptibility as a novel breeding strategy for durable and broad-spectrum resistance. Mol Breed 25: 1–12

Porebski S, Bailey LG, Baum BR (1997) Modification of a CTAB DNA extraction protocol for plants containing high polysaccharide and polyphenol components. Plant Mol Biol Report 15: 8–15

Qiang X, Liu X, Wang X, Zheng Q, Kang L, Gao X, Wei Y, Wu W, Zhao H, Shan W (2021) Susceptibility factor RTP1 negatively regulates *Phytophthora parasitica* resistance via modulating UPR regulators bZIP60 and bZIP28. Plant Physiol

Reddy ASN, Ali GS, Celesnik H, Day IS (2011) Coping with stresses: roles of calcium- and calcium/calmodulin-regulated gene expression. Plant Cell 23: 2010–2032

Schie CCNV, Takken FLW (2014) Susceptibility genes 101: how to be a good host. Annu Rev Phytopathol 52: 551–581

Schoonbeek HJ, Wang HH, Stefanato FL, Craze M, Bowden S, Wallington E, Zipfel C, Ridout CJ (2015) Arabidopsis EF-Tu receptor enhances bacterial disease resistance in transgenic wheat. New Phytol 206: 606–613

Schultheiss H, Dechert C, Kogel K-H, Hückelhoven R (2002) A small GTP-binding host protein is required for entry of powdery mildew fungus into epidermal cells of barley. Plant Physiol 128: 1447–1454

Schultheiss H, Dechert C, Kogel K-H, Hückelhoven R (2003) Functional analysis of barley RAC/ROP G-protein family members in susceptibility to the powdery mildew fungus. Plant J 36: 589–601

Schwessinger B (2017) Fundamental wheat stripe rust research in the 21st century. New Phytol 213: 1625–1631

Shimizu T, Nakano T, Takamizawa D, Desaki Y, Ishii-Minami N, Nishizawa Y, Minami E, Okada K, Yamane H, Kaku H, et al. (2010) Two LysM receptor molecules, CEBiP and OsCERK1, cooperatively regulate chitin elicitor signaling in rice. Plant J 64: 204–214

Takabatake R, Karita E, Seo S, Mitsuhara I, Kuchitsu K, Ohashi Y (2007) Pathogen-induced calmodulin isoforms in basal resistance against bacterial and fungal pathogens in tobacco. Plant Cell Physiol 48: 414–423

Tuteja N, Mahajan S (2007) Calcium signaling network in plants. Plant Signal Behav 2: 79–85

Waadt R, Schmidt LK, Lohse M, Hashimoto K, Bock R, Kudla J (2008) Multicolor bimolecular fluorescence complementation reveals simultaneous formation of alternative CBL/CIPK complexes in planta. Plant J 56: 505–516

Wang C, Huang L, Buchenauer H, Han Q, Zhang H, Kang Z (2007) Histochemical studies on the accumulation of reactive oxygen species (O^2-^ and H_2_O_2_) in the incompatible and compatible interaction of wheat - *Puccinia striiformis* f. sp *tritici*. Physiol Mol Plant Pathol 71: 230–239

Wang J, Liu X, Zhang A, Ren Y, Wu F, Wang G, Xu Y, Lei C, Zhu S, Pan T, et al. (2019) A cyclic nucleotide-gated channel mediates cytoplasmic calcium elevation and disease resistance in rice. Cell Res 29: 820–831

Wang Q, Chen B, Liu P, Zheng M, Wang Y, Cui S, Sun D, Fang X, Liu C, Lucas WJ, et al. (2009a) Calmodulin binds to extracellular sites on the plasma membrane of plant cells and elicits a rise in intracellular calcium concentration. J Biol Chem 284: 12000–12007

Wang X, Tang C, Zhang G, Li Y, Wang C, Liu B, Qu Z, Zhao J, Han Q, Huang L, et al. (2009b) cDNA-AFLP analysis reveals differential gene expression in compatible interaction of wheat challenged with *Puccinia striiformis* f. sp. *tritici*. BMC Genomics 10: 289

Wang Y, Cheng X, Shan Q, Zhang Y, Liu J, Gao C, Qiu J (2014) Simultaneous editing of three homoeoalleles in hexaploid bread wheat confers heritable resistance to powdery mildew. Nat Biotechnol 32: 947–951

Xu W, Meng Y, Surana P, Fuerst G, Nettleton D, Wise RP (2015) The knottin-like *Blufensin* family regulates genes involved in nuclear import and the secretory pathway in barley-powdery mildew interactions. Front Plant Sci 6: 409

Xu Z, Xu X, Gong Q, Li Z, Li Y, Wang S, Yang Y, Ma W, Liu L, Zhu B, et al. (2019) Engineering broad-spectrum bacterial blight resistance by simultaneously disrupting variable tale-binding elements of multiple susceptibility genes in rice. Mol Plant 12: 1434–1446

Yin C, Chen X, Wang X, Han Q, Kang Z, Hulbert SH (2009) Generation and analysis of expression sequence tags from haustoria of the wheat stripe rust fungus *Puccinia striiformis* f. sp. *tritici*. BMC Genomics 10: 626

Zhang Y, Bai Y, Wu G, Zou S, Chen Y, Gao C, Tang D (2017) Simultaneous modification of three homoeologs of *TaEDR1* by genome editing enhances powdery mildew resistance in wheat. Plant J 91: 714–724

Zhu X, Caplan J, Mamillapalli P, Czymmek K, Dinesh-Kumar SP (2010) Function of endoplasmic reticulum calcium ATPase in innate immunity-mediated programmed cell death. Embo J 29: 1007–1018

